# T-cell immunoglobulin and mucin (TIM) contributes to Hantaan virus entry into human airway epithelial cells

**DOI:** 10.1101/872317

**Authors:** Jennifer Mayor, Giulia Torriani, Gert Zimmer, Sylvia Rothenberger, Olivier Engler

## Abstract

Hantaviruses are rodent-borne haemorrhagic fever viruses leading to serious diseases. Viral attachment and entry represent the first steps in virus transmission and are promising targets for antiviral therapeutic intervention. Here we investigated receptor use in human airway epithelium of the Old and New World hantaviruses Hantaan virus (HTNV) and Andes virus (ANDV). Using a biocontained recombinant vesicular stomatitis virus pseudotype platform, we provide first evidence for a role of the cellular phosphatidylserine (PS) receptors of the T-cell immunoglobulin and mucin (TIM) in HTNV and ANDV entry. In line with previous studies, HTNV, but not ANDV, was able to use the glycosaminoglycan heparan sulfate and αvβ3 integrin as co-receptors. In sum, our studies demonstrate for the first time that hantaviruses use PS receptors and hence apoptotic mimicry to invade human airway epithelium, which may explain why these viruses can easily break the species barrier.

## INTRODUCTION

Hantaviruses are emerging negative-strand viruses associated with severe human diseases. The prototypic Hantaan virus (HTNV) and Seoul virus (SEOV) are widespread in Asia where they can cause hemorrhagic fever with renal syndrome (HFRS) with up to 15% case-fatality. The New World hantaviruses Sin Nombre (SNV) and Andes (ANDV) are associated with hantavirus cardiopulmonary syndrome (HCPS) in the Americas with up to 40% mortality. In Europe, Puumala virus (PUUV) causes *nephropathia epidemica*, a milder form of HFRS, while Dobrava-Belgrade virus (DOBV) in the Balkans is associated with the more severe form of HFRS (Jonsson, Figueiredo et al. 2010, Vaheri, Henttonen et al. 2013). The lack of a licensed vaccine and the limited therapeutic options make the development of novel antiviral strategies against hantaviruses an urgent need.

Rodents and insectivores such as shrews, moles and bats constitute the natural reservoir of hantaviruses. Hantaviruses cause asymptomatic persistent infections in their hosts, and zoonotic transmission occurs via aerosols of contaminated rodent excreta (Vaheri, Henttonen et al. 2013, Vaheri, Strandin et al. 2013). Cells of the human respiratory epithelium represent likely early targets, and after initial replication at the site of entry, hantaviruses can enter the bloodstream and disseminate systemically. In cases of severe infection, viral antigen is detectable in dendritic cells, macrophages, lymphocytes, and in microvascular endothelial cells, whose functional perturbation contributes to the fatal shock syndrome (Muranyi, Bahr et al. 2005, Vaheri, Strandin et al. 2013, Manigold and Vial 2014).

Viral attachment and entry represent the first and most fundamental steps in hantavirus zoonotic transmission and infection. Several cellular receptors have been shown to interact with glycoproteins of hantaviruses, in particular αvβ3 integrin, decay-accelerating factor (DAF) and component of the complement system (Gavrilovskaya, Shepley et al. 1998, Gavrilovskaya, Brown et al. 1999, Raymond, Gorbunova et al. 2005, Krautkramer and Zeier 2008, Popugaeva, Witkowski et al. 2012). More recently, protocadherin-1 (PCDH1) has been shown to be essential for entry of New World hantaviruses into vascular endothelial cells *in vitro* and *in vivo* (Jangra, Herbert et al. 2018). However, the receptors used by Old and New World hantaviruses in human respiratory epithelial cells are currently unknown.

Since virus-host co-evolution is driven mainly by long-term relationships of viruses with their reservoir hosts, there is no a priori selection pressure to evolve the capacity to recognize human receptors. An elegant strategy of infection that many viruses evolved consists of acquiring phosphatidylserine (PS) in their envelope during budding, and consequently be disguised as apoptotic bodies that can be engulfed by cells through clearance mechanisms (Moller-Tank and Maury 2014, Amara and Mercer 2015). Viral cell entry via this mechanism of “apoptotic mimicry” was first demonstrated by Mercer and Helenius for the poxvirus vaccinia (Mercer and Helenius 2008). Apoptotic mimicry is recognized as a major entry strategy used by a broad spectrum of viruses, including important emerging pathogens, such as Ebola (EBOV), Dengue (DENV), West Nile (WNV), and Zika (ZIKV) virus (Shimojima, Ikeda et al. 2007, Brindley, Hunt et al. 2011, Hunt, Kolokoltsov et al. 2011, Kondratowicz, Lennemann et al. 2011, Meertens, Carnec et al. 2012, Jemielity, Wang et al. 2013, Meertens, Labeau et al. 2017). The major classes of cellular PS receptors currently implicated in viral entry via apoptotic mimicry are molecules of the T-cell immunoglobulin and mucin (TIM) receptor family, in particular TIM-1 and TIM-4 and receptor tyrosine kinases of the Tyro3/Axl/Mer (TAM) family (Moller-Tank and Maury 2014, Amara and Mercer 2015).

TIM-1 and TIM-4 directly bind to PS and phosphatidylethanolamine (PE) exposed in the viral membrane via their globular head domain. Virus binding to TAM receptors requires the PS binding proteins Gas6 and protein S that are present in serum and provide a molecular bridge between TAM receptors and the virus (Fernandez-Fernandez, Bellido-Martin et al. 2008, Morizono, Xie et al. 2011, Morizono and Chen 2014, Amara and Mercer 2015). The PS receptors of the TIM and TAM families are widely expressed in many tissues, including human airway epithelial cells, vascular endothelial cells, hepatocytes, different classes of immune cells, kidney, nervous tissues, heart and skeletal muscles (Monney, Sabatos et al. 2002, Umetsu, Lee et al. 2005, Kobayashi, Karisola et al. 2007, Linger, Keating et al. 2008, Kondratowicz, Lennemann et al. 2011), conferring broad tissue tropism. In addition to their role in removal of apoptotic cells, TAM receptors play a crucial role in the negative regulation of innate immune signaling, modulating the host cell’s type interferon (IFN)-I response (Rothlin, Ghosh et al. 2007, Lemke and Rothlin 2008, Rothlin and Lemke 2010). Engagement of TAM receptors by enveloped viruses, down-regulates innate immune signaling, blunting the IFN-I response thus promoting viral replication at the post-entry level (Bhattacharyya, Zagorska et al. 2013, Moller-Tank and Maury 2014, Amara and Mercer 2015). In the present study, we investigated the possible role of TIM/TAM receptors and apoptotic mimicry in productive cell entry of the prototypic highly pathogenic Old and New World hantaviruses HTNV and ANDV into human respiratory epithelial cells. Our data provide first evidence for a role of TIM-1 in hantavirus cell entry.

## RESULTS

### Human TIM-1 promotes the entry of HTNV and ANDV pseudoviruses

Over the last years, studies have shown that human TIM-1, a protein initially implicated as a receptor for the non-enveloped hepatitis A virus, could promote infection of a range of enveloped viruses, among them filoviruses, flaviviruses, New World arenaviruses and alphaviruses (Kondratowicz, Lennemann et al. 2011, Meertens, Carnec et al. 2012, Jemielity, Wang et al. 2013, Brouillette, Phillips et al. 2018). Because TIM-1 is known to be expressed in a broad range of epithelial cells, in particular on human airway epithelium (Kondratowicz, Lennemann et al. 2011), which is likely to constitute an early cellular target for transmission of hantaviruses, we explored the possibility that TIM-1 mediates their cell entry.

A major challenge for the identification of cellular factors required for attachment and entry of hantaviruses is the strict biosafety requirements limiting work with these pathogens to BSL-3 facilities. For our study, we employed vesicular stomatitis virus (VSV)-pseudotypes we have previously characterized and validated (Torriani, Mayor et al. 2019). To generate pseudoviruses bearing the glycoproteins of the prototypic Old World hantavirus HTNV and the South American ANDV, we used recombinant VSV*∆G-Luc in which the glycoprotein (G) gene was deleted and replaced with reporter genes encoding enhanced green fluorescent protein (EGFP; indicated by an asterisk) and firefly luciferase (Luc) (Berger Rentsch and Zimmer 2011). VSV*∆G-Luc was propagated on VSV-G protein-expressing cells resulting in VSV-G *trans*-complemented VSV*∆G-Luc(VSV-G) particles. To produce VSV*∆G-Luc(HTNV-G) and VSV*ΔG-Luc(ANDV-G), HEK293F cells were transfected with a plasmid expressing HTNV or ANDV glycoproteins, infected with VSV*∆G-Luc(VSV-G) particles and incubated in the presence of the neutralizing anti-VSVG monoclonal antibody I1 (Holland, de la Torre et al. 1989) to lower background (Fig. 1A). Using a similar approach, we generated pseudoviruses for EBOV and Lassa virus (LASV). The resulting “VSV-pseudotypes” are replication competent, but unable to propagate, making them suitable for work under BSL-2 conditions.

**FIGURE 1.**
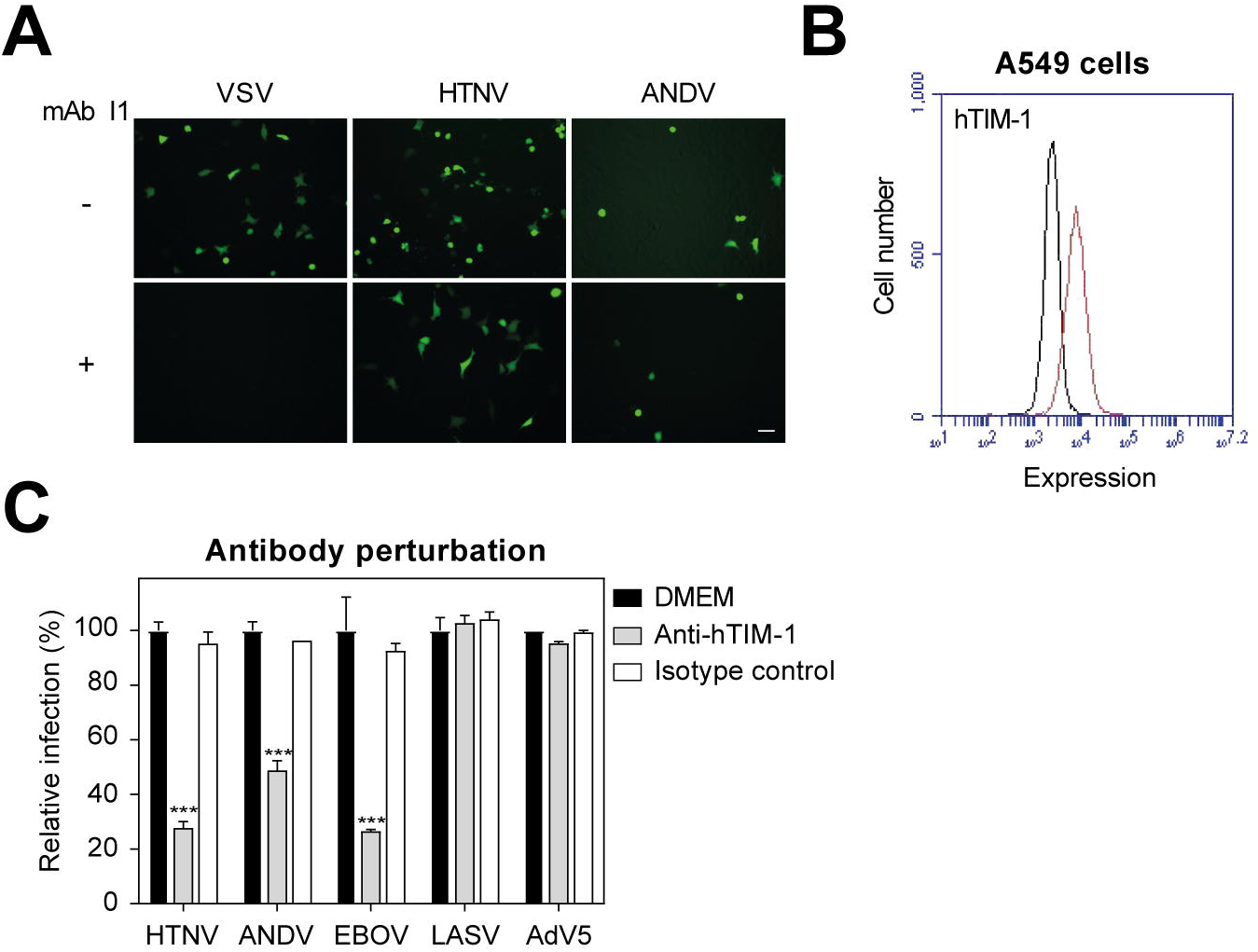
hTIM-1 can mediate productive entry of HTNV-G and ANDV-G. **(A)** Specificity to neutralizing activity of anti-VSVG antibody. A549 cells were infected with VSV*ΔG-Luc(VSV-G), VSV*ΔG-Luc(HTNV-G), VSV*ΔG-Luc(ANDV-G), in the presence or absence of the VSV-G neutralizing antibody I1 (mAb I1). **(B)** Detection of hTIM-1 on A549 cells by flow cytometry. Fixed non-permeabilised cells were stained with goat anti-TIM-1 pAb. Black line represents secondary antibody only, and red line represents primary and secondary antibodies. **(C)** Blocking of hTIM-1-mediated infection by antibody perturbation. A549 cells were pre-treated for 1 h at 4 °C with 30 nM of goat anti-hTIM-1 pAb and control goat IgG in the absence of serum. Cells were then infected with VSV*ΔG-Luc(HTNV-G), VSV*ΔG-Luc(ANDV-G), VSV*ΔG-Luc(EBOV-G), VSV*ΔG-Luc(LASV-G) and AdV5 and shifted to 37 °C for 90 min. Cells were washed with medium supplemented with 20 mM ammonium chloride, followed by 16 h of incubation in the presence of lysosomotropic agent. Infection was quantified by counting EGFP-positive infected cells. Data are normalized to the control DMEM and displayed as percentage of infection related to the control. Data are means + SD (n=3) with p-value ***: p ≤ 0.001.

As hantaviruses are transmitted via aerosols (Vaheri, Henttonen et al. 2013, Vaheri, Strandin et al. 2013), we used A549 human lung epithelial cells which are highly susceptible to infection with HTNV and ANDV pseudoviruses as a suitable cell model. Using flow cytometry, we detected high levels of endogenous TIM-1 on the surface of A549 cells (Fig. 1B), in line with the known expression pattern of TIM-1 in A549 cells and *in vivo* (Meertens, Carnec et al. 2012). To assess the role of endogenous hTIM-1 in cell entry of hantaviruses, we tested the ability of anti-TIM-1 antibody to block infection. To do this, we treated A549 cells with goat anti-TIM-1 polyclonal antibody or normal goat IgG as control, and subsequently infected the cells with either VSV*∆G-Luc(HTNV-G) or VSV*ΔG-Luc(ANDV-G) in presence of the antibodies. As shown in Fig. 1C, hTIM-1 blocking antibody significantly reduced the entry of VSV*∆G-Luc(HTNV-G) and VSV*ΔG-Luc(ANDV-G). The antibody perturbation experiment also significantly blocked the entry of VSV*∆G-Luc(EBOV-G), used as a control, in line with previous findings (Kondratowicz, Lennemann et al. 2011). In contrast, the antibody treatment did not perturb the entry of VSV*∆G-Luc(LASV-G), as expected, since A549 cells express large amounts of functional α-Dystroglycan (α-DG) (Brouillette, Phillips et al. 2018), and had no impact on the entry of a non-enveloped virus such as recombinant adenovirus 5 (AdV5).

To examine whether hTIM-1 enhances hantavirus infection, we took advantage of the human embryonic cell line HEK293 which expresses low detectable levels of hTIM-1/-4 (Kondratowicz, Lennemann et al. 2011, Meertens, Carnec et al. 2012, Brouillette, Phillips et al. 2018) and generated cell lines overexpressing hTIM-1 (Fig. 2A). Analysis by direct fluorescence microscopy revealed that the entry of VSV*∆G-Luc(HTNV-G) and VSV*ΔG-Luc(ANDV-G) was facilitated in cells overexpressing hTIM-1 compared to parental HEK293 cells (Fig. 2B). Parental and hTIM-1 expressing HEK293 cells were infected by a panel of pseudoviruses bearing the glycoproteins of HTNV, ANDV, EBOV and LASV, as well as recombinant AdV5. Our results show that the entry of VSV*∆G-Luc(HTNV-G) and VSV*∆G-Luc(ANDV-G) was strongly enhanced in cells ectopically expressing hTIM-1 compared to the parental cells (Fig. 2C). Remarkably, the entry of VSV*∆G-Luc(HTNV-G) was increased to the same level as VSV*∆G-Luc(EBOV-G), which was taken as a control. Overexpression of hTIM-1 had no influence on the entry of VSV*∆G-Luc(LASV-G) and AdV5, in concordance with the antibody perturbation experiments in A549 cells (Fig. 1C). Quantifications upon infections from 50 to 400 infectious unit (IU)/well revealed that ectopic expression of hTIM-1 increased entry of VSV*∆G-Luc(HTNV-G) and VSV*∆G-Luc(ANDV-G) up to 100-fold and 6-fold, respectively (Fig. 2D). Taken together the data show that hTIM-1 contributes to the cell entry of hantaviruses with possible strain-specific differences.

**FIGURE 2.**
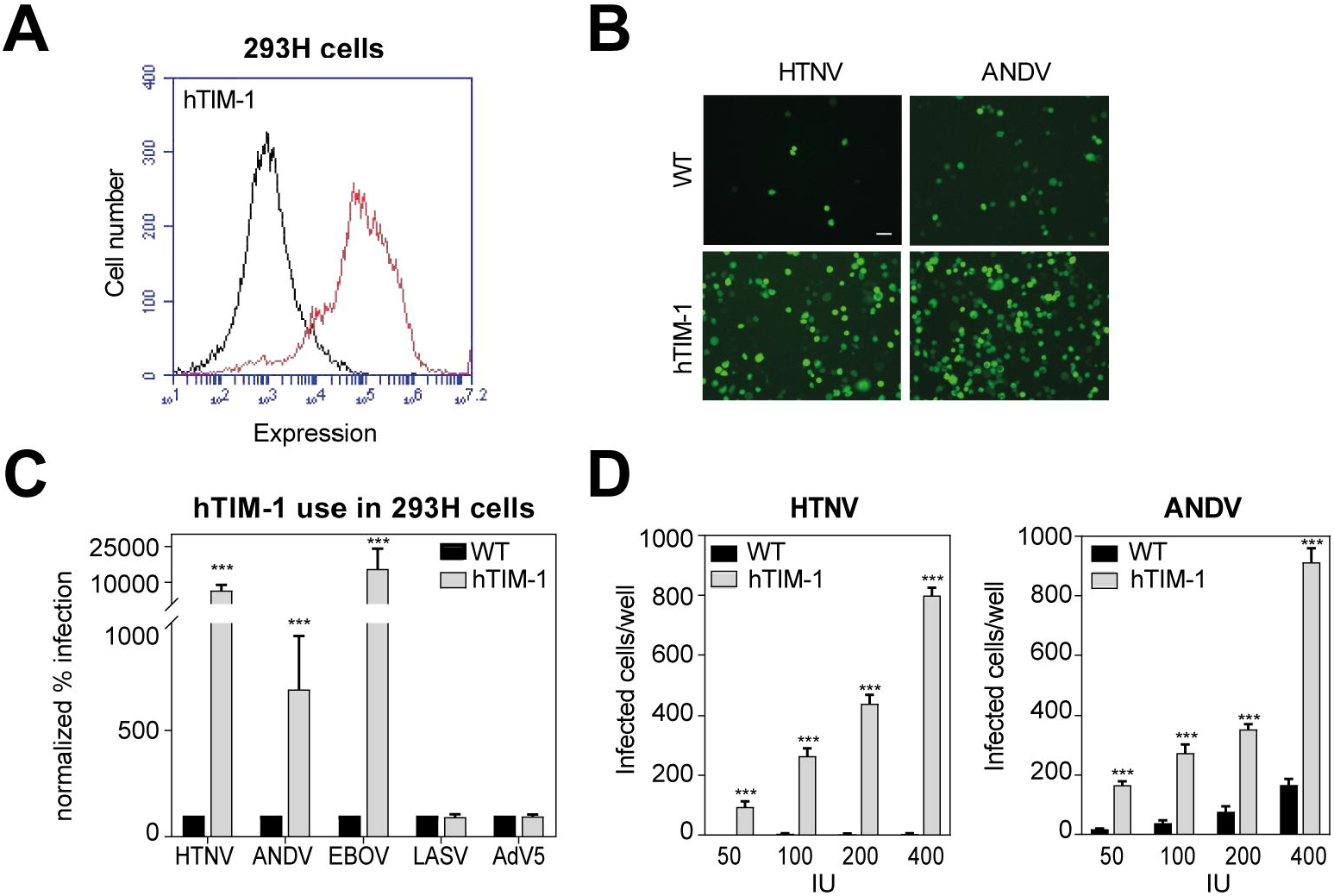
hTIM-1 promotes infection of HTNV-G and ANDV-G. **(A)** Detection of hTIM-1 on 293H cells by flow cytometry. Expression levels of hTIM-1 (red line) were assessed as in Fig. 1B. **(B)** Example of infection of VSV*ΔG-Luc(HTNV-G) and VSV*ΔG-Luc(ANDV-G) in 293H cells overexpressing hTIM-1 visualized by EGFP fluorescence of infected cells. **(C)** hTIM-1 use in 293H cells. Monolayers of 293H and derivate transfected cells were infected with VSV*ΔG-Luc(HTNV-G), VSV*ΔG-Luc(ANDV-G), VSV*ΔG-Luc(EBOV-G), VSV*ΔG-Luc(LASV-G) and AdV5-GFP at 200 IU. Infection was quantified by counting EGFP-positive infected cells. Data are normalized to the control 293H WT cells and displayed as normalized percentage of infection related to the control. **(D)** Overexpression of hTIM-1 in 293H cells. Monolayers of 293H and derivate transfected cells were infected with VSV*ΔG-Luc(HTNV-G) and VSV*ΔG-Luc(ANDV-G) at the indicated IU. Infection was quantified by counting EGFP-positive infected cells or measuring the luciferase. **(C-E)** Data are means + SD (n=3) with p-value ***: p ≤ 0.001.

### Viral usage of the PS receptor hTIM-4 parallels that of hTIM-1

Studies identified TIM-4 as a potent entry enhancer of a wide range of viruses including DENV and WNV (Meertens, Carnec et al. 2012, Jemielity, Wang et al. 2013). Whereas hTIM-1 is highly expressed in epithelial cells, hTIM-4 expression is restricted to dendritic cells and macrophages (Kobayashi, Karisola et al. 2007, Miyanishi, Tada et al. 2007). In order to test whether hTIM-4 could also promote hantavirus entry, we transiently expressed hTIM-4 in HEK293 cells (Fig. 3A). Parental and hTIM-4 expressing cells were incubated in presence of a panel of pseudoviruses as above, and the infection followed by measuring luciferase activity at 50 to 200 IU/well (Fig. 3B and 3C). Our data show that the viral usage of hTIM-4 parallels that of hTIM-1, although at a lower efficiency. These results further open the possibility that hTIM-4 mediates the entry of hantaviruses in macrophages and dendritic cells.

**FIGURE 3.**
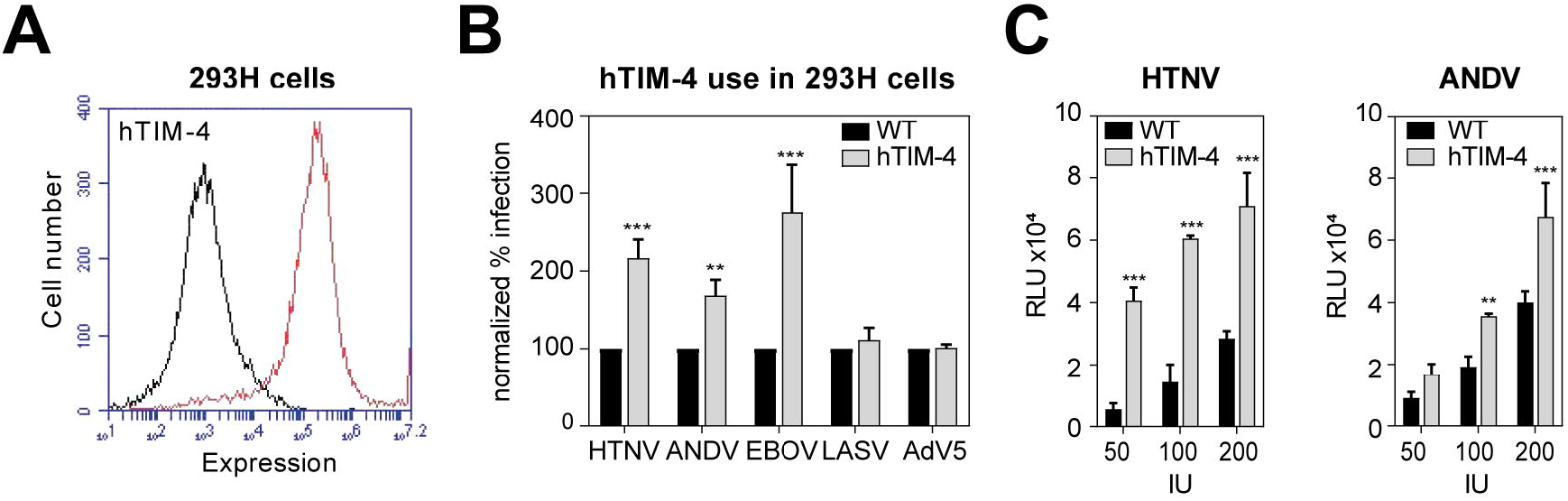
Viral usage of the PS receptor hTIM-4 parallels that of hTIM-1. **(A)** Detection of hTIM-4 on 293H cells by flow cytometry. Surface levels of hTIM-4 on 293H and transfected cells were evaluated by flow cytometry using the goat anti-hTIM-4 pAb (red line). **(B)** hTIM-4 use in 293H cells. Infection of 293H and derivate TIM-4 cells was assessed as in Fig. 2C. Infection was quantified by counting EGFP-positive infected cells or measuring the luciferase. **(C)** Overexpression of hTIM-4 in 293H cells. Infection of confluent monolayers of 293H and derivate transfected cells was similarly assessed as in Fig. 2D. Infection was detected by luciferase assay. **(B-C)** Data are means + SD (n=3) with p-value **: p ≤ 0.01; ***: p ≤ 0.001.

### The receptor-type tyrosine kinase Axl promotes the entry of ANDV-G

Whereas TIM-1 and TIM-4 directly bind to PS and PE exposed in the viral membrane, virus binding to TAM receptors requires the PS binding proteins Gas6 and protein S that are present in the serum and serve as molecular bridge between TAM receptors and the virus (Fernandez-Fernandez, Bellido-Martin et al. 2008, Morizono, Xie et al. 2011, Morizono and Chen 2014, Amara and Mercer 2015). The tyrosine receptor kinase Axl is one of the three members of the TAM (Tyro3, Axl, Mer) protein family. Axl is ubiquitously expressed in human tissues and is implicated in multiple functions, including cell survival, proliferation, migration, adhesion, clearance of apoptotic material and immune regulation (Linger, Keating et al. 2008). In addition, Axl was shown to enhance the entry of EBOV (Shimojima, Takada et al. 2006, Shimojima, Ikeda et al. 2007, Brindley, Hunt et al. 2011), and is the primary ZIKV entry co-factor (Meertens, Labeau et al. 2017, Richard, Shim et al. 2017). To examine the requirement for Axl, we treated A549 human lung epithelial cells with polyclonal antibodies directed against Axl ectodomain that have been previously shown to inhibit EBOV entry (Shimojima, Takada et al. 2006, Brindley, Hunt et al. 2011). A549 cells do express Axl, as detected by flow cytometry (Fig. 4A). The cells were treated with the Axl antisera or normal goat IgG as control, and subsequently infected with either VSV*∆G-Luc(HTNV-G) or VSV*ΔG-Luc(ANDV-G). Our analysis showed that Axl antibody significantly reduced VSV*∆G-Luc(ANDV-G) entry, but not VSV*∆G-Luc(HTNV-G), indicating a strain-specific usage of this receptor in A549 cells (Fig. 4B). Axl antisera inhibited VSV*∆G-Luc(EBOV-G) entry, in line with previous reports (Shimojima, Takada et al. 2006, Shimojima, Ikeda et al. 2007, Brindley, Hunt et al. 2011). Entry of AdV5 was not perturbed, as expected. Moreover, combination of both antibodies demonstrated a higher inhibition, and an additive effect of VSV*∆G-Luc(ANDV-G) and VSV*∆G-Luc(EBOV-G) entry. Our results suggest that Axl and TIM-1 are differently used by HTNV and ANDV pseudoviruses to enter into host cells. Activation of Axl regulates a number of signal transduction pathways and signaling may be one important component of Axl-dependent virus entry, as shown by the requirement of Axl cytoplasmic tail residues for EBOV entry (Shimojima, Ikeda et al. 2007). To evaluate the impact of Axl-dependent signaling we used the novel small-molecule Axl tyrosine kinase inhibitor R428, which shows high selectivity and fast drug action (Holland, Pan et al. 2010, Myers, Brunton et al. 2016). A549 cells were pre-treated with R428 for 30 min at 37°C, followed by an incubation with the pseudoviruses in presence of the drug. VSV*∆G-Luc(EBOV-G) and AdV5 were used as positive and negative controls, respectively. After 90 min the drug was washed out to minimize the duration of drug exposure and unwanted off-target effects and the cells further incubated in medium supplemented with ammonium chloride to block further entry via low pH-triggered membrane fusion. The number of infected cells was evaluated after 16 h by counting the number of EGFP-positive infected cells. The Axl inhibitor R428 reduced the entry of VSV*∆G-Luc(ANDV-G) and VSV*∆G-Luc(EBOV-G) pseudoviruses in a dose-dependent manner (Fig. 4C). In contrast, the inhibition of VSV*∆G-Luc(HTNV-G) was less pronounced and not dose-dependent. The R428 kinase inhibitor applied under the assay conditions demonstrated no cytotoxic effect, as shown by a measure of the cell viability (Fig. 4C).

**FIGURE 4.**
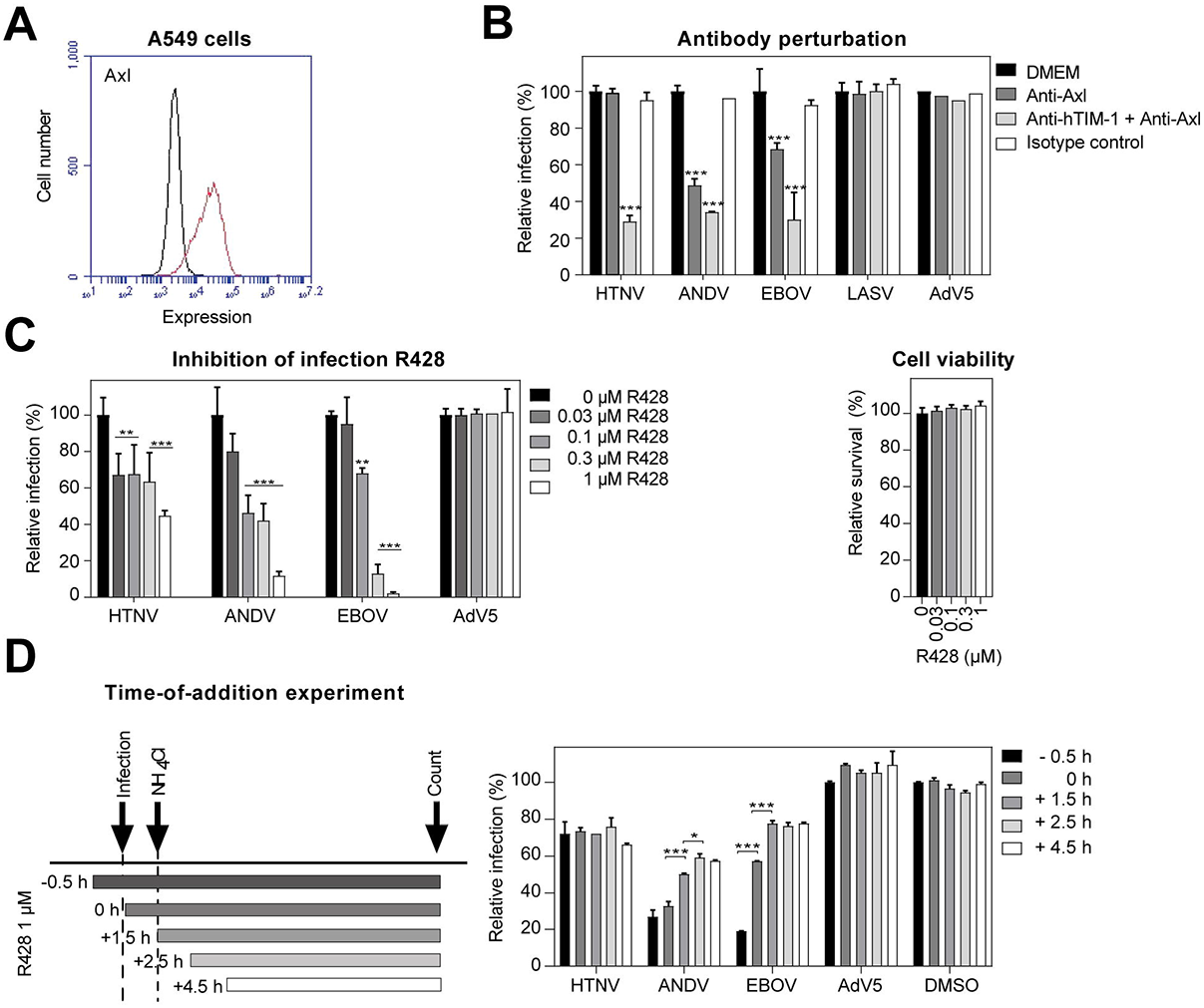
Differential role of Axl in cell entry of HTNV-G and ANDV-G pseudoviruses. **(A)** Detection of Axl on A549 cells. Expression level of Axl (red line) was assessed as in Fig. 1B. **(B)** Axl contributes to entry of ANDV-G but not HTNV-G pseudoviruses into A549 cells. A549 cells were blocked with 30 nM of goat anti-Axl pAb, goat anti-hTIM-1 pAb and control goat IgG for 1 h in the cold. Cells were then infected with indicated pseudoviruses and shifted to 37 °C. After 90 min, complete medium containing 20 mM ammonium chloride was added, followed by 16 h of incubation. Infection levels were assessed by counting EGFP-positive infected cells. **(C)** Inhibition of Axl RTK differentially affects entry of ANDV and HTNV pseudotypes into A549 cells. A549 cells were pre-treated with the Axl tyrosine kinase inhibitor R428 at increasing concentrations for 30 min, and infected with VSV*ΔG-Luc(HTNV-G), VSV*ΔG-Luc(ANDV-G), VSV*ΔG-Luc(EBOV-G) and AdV5-GFP at 200 IU/well for 90 min. Complete medium containing 20 mM ammonium chloride was added, followed by 16 h of incubation and infection levels were assessed by counting EGFP-positive infected cells. R428 shows no overt toxicity up to 1 uM. Cell viability was assessed by the CellTiter-Glo® assay, based on the antibody perturbation experiment settings. Data are means + SD (n=3) of Relative Light Units (RLU). **(D)** Time-of-addition experiment. A549 cells were infected with VSV*ΔG-Luc(HTNV-G), VSV*ΔG-Luc(ANDV-G), VSV*ΔG-Luc(EBOV-G) and AdV5-GFP at 200 IU/well with 1 uM R428 added at different time points pre-or post-infection. Pseudoviruses were added at time point 0, followed by washing and addition of ammonium chloride after 1.5, 2.5 and 4.5 h. Ammonium chloride was kept throughout the experiment and infection levels were assessed by counting EGFP-positive infected cells. Cell viability was determined by CellTiter-Glo® assay. **(B-D)** Data are means + SD (n=3) with p-value *: p ≤ 0.05; **: p ≤ 0.01; ***: p ≤ 0.001.

To delineate the impact of the drug on viral entry and post-entry steps of infection we performed a “time-of-addition” experiment. To do this, the inhibitor R428 was added at a concentration of 1 μM at different time points, before or after viral infection and maintained during the time-course of the experiment (Fig. 4D). We have previously examined the entry kinetics of our pseudoviruses and found that endosomal escape of VSV*∆G-Luc(HTNV-G) and VSV*∆G-Luc(ANDV-G) pseudotypes occurred with half times of 28 min and 42 min, respectively (Torriani, Mayor et al. 2019). Concordant with the wash-out experiment, R428 inhibited the entry of VSV*∆G-Luc(ANDV-G) and VSV*∆G-Luc(EBOV-G) pseudoviruses (Fig. 4D). In addition to the effect on viral entry, an additional inhibitory effect was measured when R428 was added 1.5 to 4.5 h after infection, indicating that R428 has also an impact on later steps of the viral cycle. These post-entry effects of TAM receptors signaling may be linked to the negative regulation of innate signaling by modulating the host cell’s type I interferon (IFN-I) response (Rothlin, Ghosh et al. 2007, Lemke and Rothlin 2008, Rothlin and Lemke 2010).

### TIM-1, αvβ3 integrin and heparan sulfate contribute to HTNV-G entry

Previous studies have demonstrated that the entry of pathogenic hantaviruses is facilitated by α_V_β_3_ integrin in human umbilical vein endothelial cells (HUVEC) and Vero E6 cells (Gavrilovskaya, Shepley et al. 1998, Gavrilovskaya, Brown et al. 1999). The human epithelial cell line A549 expresses α_V_β_3_ integrin at the cell surface, as shown by flow cytometry (Fig. 5A). To determine whether α_V_β_3_ integrin is involved in hantavirus entry, we pre-treated A549 cells with mouse monoclonal antibody anti-α_V_β_3_, or isotype control. The cells were then infected with pseudoviruses bearing the glycoproteins of HTNV, ANDV or EBOV in presence of the antibodies. After 90 min, the inoculum was removed, and the cells further incubated in presence of ammonium chloride to block further entry via low pH-triggered membrane fusion. Whereas the α_V_β_3_ integrin antibody reduced the entry of VSV*∆G-Luc(HTNV-G), it did not reduced significantly the entry of VSV*∆G-Luc(ANDV-G) (Fig. 5B). As expected, the anti-α_V_β_3_ antibody did not block the entry of VSV*∆G-Luc(EBOV-G). We have shown that the PS receptor TIM-1 greatly contributes to the entry of VSV*∆G-Luc(HTNV-G) and VSV*∆G-Luc(EBOV-G) in A549 cells (Fig. 1C and Fig. 2C, 2D). In order to evaluate the contribution of both TIM-1 and α_V_β_3_ integrin, we performed antibody-blocking experiments, treating A549 cells with either goat anti-human TIM-1 polyclonal antibody or mouse monoclonal antibody anti-α_V_β3 integrin, or a combination of both (Fig. 5C). Although anti-TIM-1 and anti-α_V_β_3_ integrin antibodies reduced the entry of VSV*∆G-Luc(HTNV-G), we did not observe an additive effect when both type of antibodies are combined.

**FIGURE 5.**
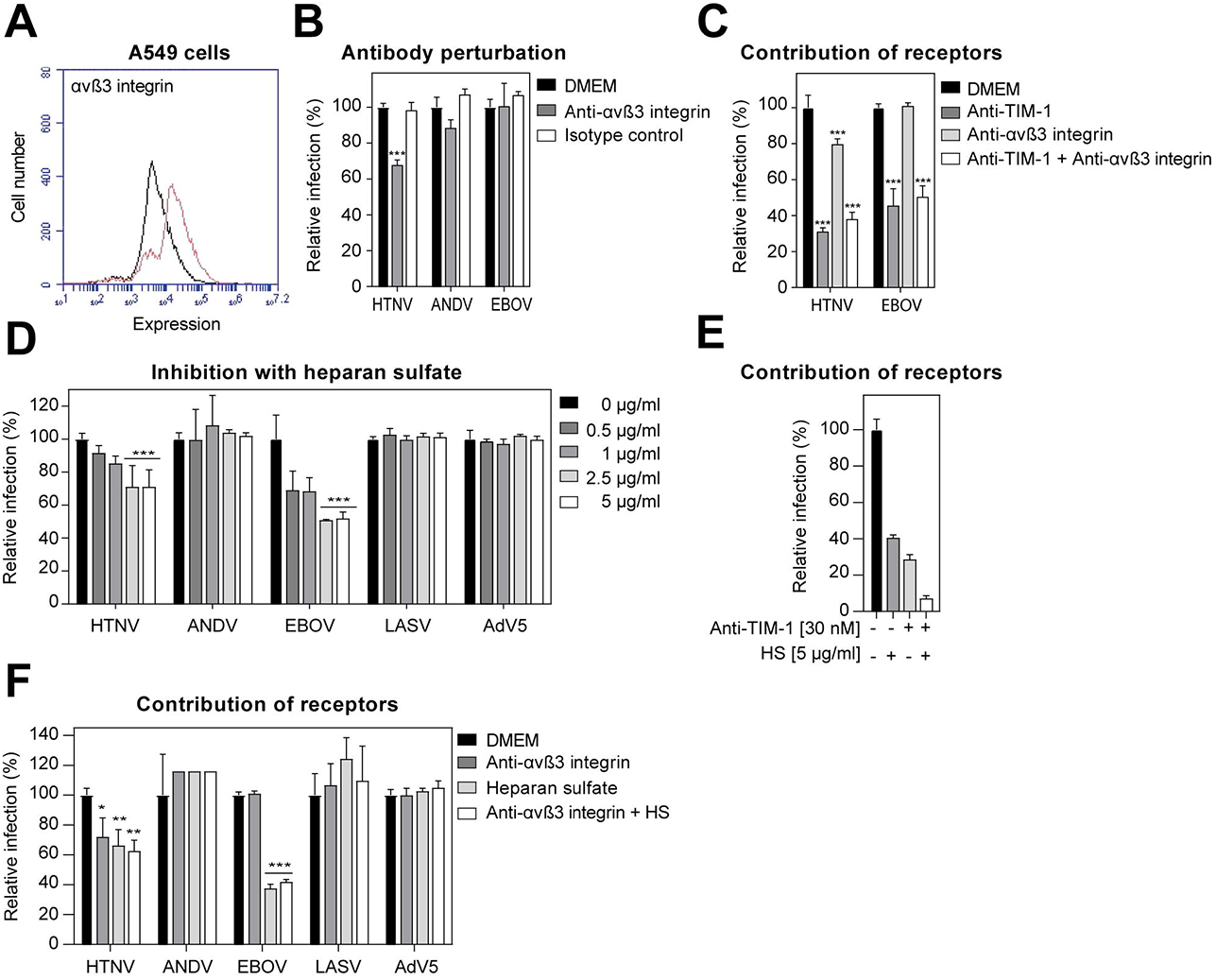
TIM-1, αvβ3 integrin, and heparan sulfate can contribute to cell entry of HTNV. **(A)** Expression of αvβ3 integrin in A549 cells. Live non-permeabilised cells were stained with the mouse anti-αvβ3 integrin mAb (red line). **(B)** Anti-αvβ3 integrin Ab reduces infection of VSV*ΔG-Luc(HTNV-G) but not VSV*ΔG-Luc(ANDV-G) into A549 cells. A549 cells were pre-treated for 1 h at 4 °C with mouse anti-αvβ3 mAb and a control mouse at 30 nM. Cells were then infected with VSV*ΔG-Luc(HTNV-G), VSV*ΔG-Luc(ANDV-G) and VSV*ΔG-Luc(EBOV-G), and shifted to 37 °C. Infection was quantified by counting EGFP-positive infected cells per well. **(C)** Relative contribution of αvβ3 integrin and hTIM-1. A549 cells were pre-treated with anti-αvβ3 and anti-hTIM-1 antibodies at 30 nM. The experiment settings were identical to the antibody perturbation assay as in panel (B). **(D)** Heparan sulfate reduces infection of HTNV but not ANDV pseudotypes into A549 cells. VSV*ΔG-Luc(HTNV-G), VSV*ΔG-Luc(ANDV-G), VSV*ΔG-Luc(EBOV-G), VSV*ΔG-Luc(LASV-G) and AdV5-GFP were pre-incubated with soluble heparan sulfate at various concentrations and used to infect monolayers of A549 cells. Infection levels were assessed by counting EGFP-positive infected cells. **(E)** Heparan sulfate and anti-hTIM-1 block HTNV-G entry. A549 cells were pre-treated for 1 h at 4 °C with the polyclonal Ab goat anti-hTIM-1 at 30 nM. Meantime, VSV*ΔG-Luc(HTNV-G), was pre-incubated with heparan sulfate (HS) at 5 µg/ml. Cells were then infected with pre-treated VSV*ΔG-Luc(HTNV-G) and shifted to 37 °C. Infection was quantified 16 h post-infection by counting EGFP-positive infected cells. **(F)** Relative contribution of αvβ3 integrin and heparan sulfate. Pre-treatment of cells with anti-αvβ3 mAb (30 nM), viruses with 5 µg/ml of heparan sulfate (HS) and infections were assessed as in Fig. 5E. Infection was quantified 16 h post-infection by counting EGFP-positive infected cells. **(B-F)** Data are means + SD (n=3) with p-value *: p ≤ 0.05; **: p ≤ 0.01; ***: p ≤ 0.001.

In addition to the PS receptors of the TIM and TAM families, heparan sulfate proteoglycan (HSPG), one of major negatively charged transmembrane protein-linking glycosaminoglycans, is involved in binding of a broad range of viruses, including DENV (Morizono and Chen 2014), EBOV (Salvador, Sexton et al. 2013, O’Hearn, Wang et al. 2015, Tamhankar, Gerhardt et al. 2018) and hepatitis C virus (Barth, Schafer et al. 2003). HSPG is expressed by almost all cells. We tested whether the glycoprotein of hantaviruses could interact with HSPG by pre-treating a panel of pseudoviruses, as well as AdV5, with soluble heparan sulfate (HS). A549 cells were inoculated with the mixtures for 90 min. The inoculum was then removed, and the cells further incubated in presence of ammonium chloride to block secondary infection via low pH-triggered membrane fusion (Fig. 5D). Entry of both EBOV and HTNV pseudoviruses was inhibited in the presence of HS in a dose-dependent manner, whereas HS did not impact the entry of VSV*∆G-Luc(ANDV-G), VSV*∆G-Luc(LASV-G) or AdV5. The HS dependence of HTNV, but not of ANDV is in line with previous findings using a genetic approach (Riblett, Blomen et al. 2016). We further combined HS and TIM-1 blocking antibody (Fig. 5E). Our results show an additive effect of HS and anti-TIM-1 blocking antibody and a drastic reduction of VSV*∆G-Luc(HTNV-G) cell entry when the two compounds are combined.

As shown above, heparan sulfate and α_V_β_3_ integrin significantly affected VSV*∆G-Luc(HTNV-G) entry when applied separately. However, we did not observe a significant additive effect when both anti-α_V_β_3_ integrin and soluble heparan sulfate are combined (Fig. 5F).

### Duramycin is a potent drug against pathogenic HTNV

PE is also a ligand for PS receptors and plays an important role in apoptotic clearance and entry of many pathogenic viruses (Richard, Zhang et al. 2015). We measured the sensitivity of our pseudoviruses to the exposure to duramycin, a lantibiotic agent that binds PE (Iwamoto, Hayakawa et al. 2007, Richard, Zhang et al. 2015). Pseudoviruses bearing the glycoproteins of HTNV, ANDV, EBOV or LASV, as well as recombinant AdV5 were pre-incubated with increasing concentrations of duramycin and subsequently incubated in presence of A549 cells. Duramycin drastically decreased cell entry of hantaviruses, with more than 90% reduction for both VSV*∆G-Luc(HTNV-G) and VSV*∆G-Luc(ANDV-G) at concentration of 0.2 μM (Fig. 6A). As expected, cell infection by AdV5 used as control was unaffected by the treatment, even at the highest concentration of 1 μM duramycin. The duramycin-mediated inhibition was further evaluated with authentic infectious HTNV in BSL-3 containment (Fig. 6B). HTNV particles were incubated with various concentration of duramycin for 30 min, followed by infection of Vero E6 cells at 37°C for 1 h. Cells were washed once and incubated with medium containing appropriate duramycin concentration for five days. We observed 88% of inhibition at 1 μM duramycin concentration. The duramycin applied under the assay conditions demonstrated no cytotoxic effect, as shown by a measure of the cell viability.

**FIGURE 6.**
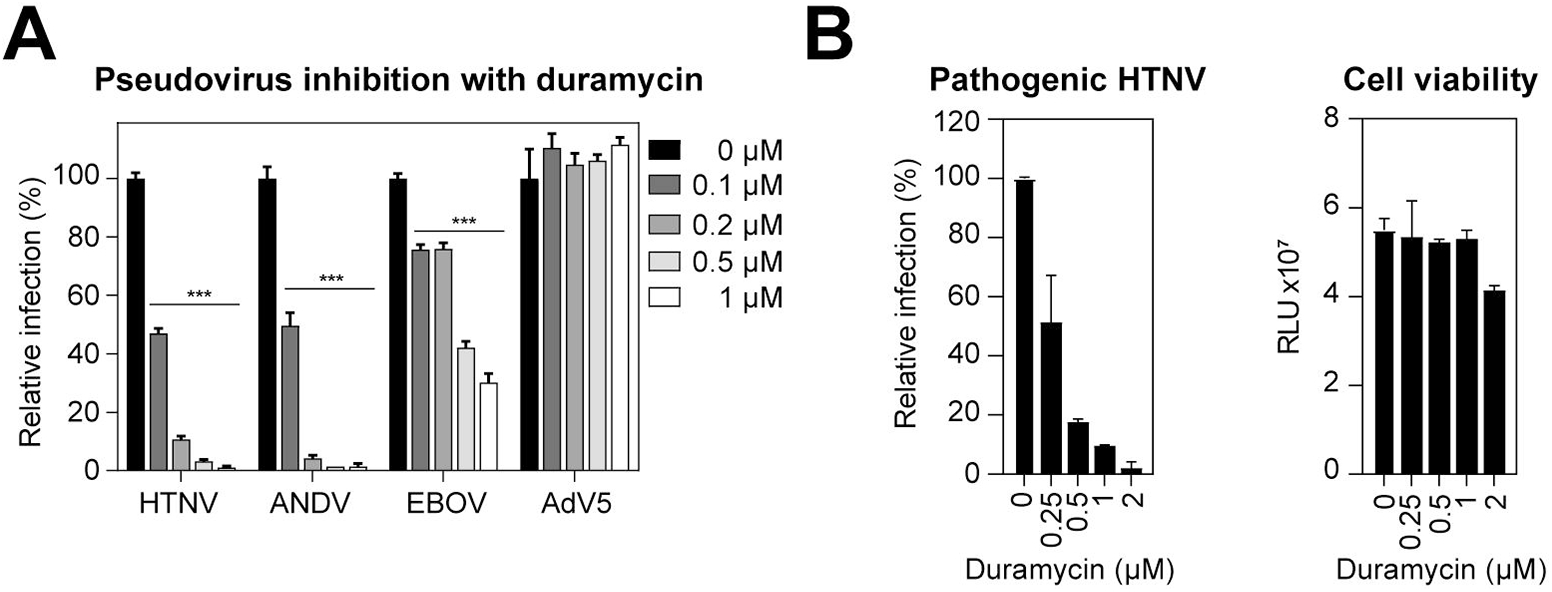
Duramycin inhibits HTNV infection. **(A)** Inhibition of HTNV-G and ANDV-G infection in A549 cells with duramycin. VSV*ΔG-Luc(HTNV-G), VSV*ΔG-Luc(ANDV-G), VSV*ΔG-Luc(EBOV-G) and AdV5-GFP were pre-incubated with duramycin at the indicated concentrations for 1 h at RT and used to infect A549 cells. After 90 min, complete medium containing 20 mM ammonium chloride was added, followed by 16 h of incubation in the presence of lysosomotropic agent. Infection levels were assessed by counting EGFP-positive infected cells. Data are means + SD (n=3) with p-value ***: p ≤ 0.001. **(B)** Inhibition of HTNV entry into Vero E6 cells with duramycin. HTNV was pre-treated for 30 min at RT with increasing concentrations of duramycin. Cells were subsequently infected for 1 h at 37°C, washed once with medium and incubated with medium containing appropriate duramycin concentrations for 5 days. Infection levels were assessed by qPCR. Cell viability was determined by CellTiter-Glo® assay. Data are means + SD (n=2).

## DISCUSSION

The emergence of novel highly pathogenic viruses represents an important problem for public health. Climate changes, almost unrestricted global trade, and increasing urbanization promote the emergence of novel human pathogenic viruses. Most newly emerging pathogenic viruses are of zoonotic origin and reservoir to human transmission is crucial for spillover into human population. Through their diversity, global distribution and capacity to cause severe disease in human, hantaviruses represent an important model for the study of zoonotic transmission via aerosols.

Receptor binding and entry are the first steps of a viral infection and are decisive for host-range, tissue tropism, and disease potential of the virus. The capacity of a zoonotic virus to break the species barrier depends on its ability to use receptor(s) in the new species allowing productive infection. Here we provide first evidence that HTNV and ANDV hijack the phosphatidylserine receptor TIM-1 to enter human epithelial cells. To circumvent biosafety restrictions, we used a biocontained recombinant VSV pseudotype platform. In the past years, VSV-derived pseudoviruses were used in multiple studies to evaluate entry factors for hantaviruses, filoviruses and arenavirus among others. The studies demonstrated a very close correlation of pseudovirions with the pathogenic viruses (Kondratowicz, Lennemann et al. 2011, Jemielity, Wang et al. 2013, Moller-Tank, Kondratowicz et al. 2013, Moller-Tank and Maury 2014, Kuroda, Fujikura et al. 2015, Riblett, Blomen et al. 2016, Brouillette, Phillips et al. 2018, Jangra, Herbert et al. 2018). Entry of HTNV and ANDV into A549 lung epithelial cells was significantly reduced in presence of anti-hTIM-1 blocking antibody (Fig. 1). Conversely, overexpression of hTIM-1 in HEK293 cells, which express low levels of hTIM-1 induces a massive increase in the uptake of HTNV and EBOV pseudovirions, and a more modest increase in the uptake of ANDV pseudovirions (Fig. 2). TIM-1 is expressed on cilia of human airway epithelia (Kondratowicz, Lennemann et al. 2011). Thereby cells of the human respiratory epithelium may represent early targets for the viruses displaying a dependency on TIM-1 for entry, such as HTNV and EBOV. This hypothesis is currently explored in the high containment laboratories at Spiez Laboratory using organotypic cultures of human airways.

Our data show that the viral usage of hTIM-4 parallels that of hTIM-1, although at a lower efficiency (Fig. 3). TIM-4 is predominantly expressed in macrophages and subsets of dendritic cells (Meyers, Chakravarti et al. 2005, Kobayashi, Karisola et al. 2007, Miyanishi, Tada et al. 2007). These results further open the possibility that hTIM-4 mediates the entry of hantaviruses in macrophages and dendritic cells.

We found by antibody perturbation experiments, as well as use of the R428 kinase inhibitor that the tyrosine receptor kinase Axl, a member of the TAM (Tyro3, Axl, Mer) protein family could promote entry of ANDV, but not HTNV (Fig. 4). This differential receptor use by these two viruses may contribute to differences in tropism and pathogenesis. Axl is ubiquitously expressed in human tissues and is implicated in multiple functions, including cell survival, proliferation, migration, adhesion, clearance of apoptotic material and immune regulation (Linger, Keating et al. 2008). In the lungs, the TAM (Tyro3, Axl, Mer) tyrosine kinase receptors mediate apoptotic cell uptake by phagocytes which is critical for lung homeostasis. TAM receptors display a strong cell-specific pattern and undergoes tight regulations. In particular, Axl expression is restricted to airway macrophages and is not detected on peripheral blood monocytes or monocyte-derived macrophages (Grabiec, Denny et al. 2017). As we observe that Axl can contribute to the entry of ANDV, lung macrophages could constitute early targets, a possibility that could be assessed using pulmonary macrophages isolated after bronchoalveolar lavage (BAL) as cell model.

Using antibody perturbation, we showed that the previously identified hantavirus receptor αvβ3 integrin (Gavrilovskaya, Shepley et al. 1998, Gavrilovskaya, Brown et al. 1999) contributes to HTNV entry in A549 cells (Fig. 5). We also found that HSPG was used by HTNV, but not ANDV in A549 cells, a finding that is line with a previous report (Riblett, Blomen et al. 2016). However, our results suggest a more important contribution of the TIM/TAM receptors compared to integrin and HSPG for entry of hantaviruses in human epithelial cells.

Recently, PCDH1 has been shown to be essential for entry of ANDV into vascular endothelial cells (Jangra, Herbert et al. 2018). It remains to be determined whether PCDH1 plays a role in human airway epithelial cells, in addition to TIM-1 and Axl.

The viral entry into host cells is mainly determined by the interaction between cellular receptors and proteins present at the surface of the enveloped virus. Based on previous studies on DENV and EBOV (Meertens, Carnec et al. 2012, Kuroda, Fujikura et al. 2015, Richard, Zhang et al. 2015), we expect duramycin to inhibit authentic hantaviruses. Indeed we showed that duramycin is a potent antiviral against pathogenic HTNV with 88% of viral infection decrease (Fig. 6). PS and PE were shown to interact with TIM-1/-4 (Meyers, Chakravarti et al. 2005, Kobayashi, Karisola et al. 2007, Miyanishi, Tada et al. 2007), however we observed that the duramycin-mediated inhibition was greater that the contribution of TIM receptors, suggesting the use of additional cellular factors. CD300a, which contributes to DENV infection (Carnec, Meertens et al. 2016) is a possible candidate. Its role in hantaviruses entry into human airway epithelial cells should be further investigated.

In sum, our studies provide first evidence for a role of TIM-1 and expand understanding on hantaviruses entry into human airway epithelial cells. A better understanding of the mechanisms of virus attachment and entry is essential to develop therapeutic strategies to fight against hantaviruses. Identification of TIM proteins as receptors give new insights into the viral transmission between natural reservoirs and human host and can lead moreover to new avenues for the development of drugs.

## MATERIALS AND METHODS

### Plasmids, antibodies, and reagents

pWRG/HTNV-M (Hooper, Custer et al. 2001) was kindly provided by Connie. S. Schmaljohn (U.S. Army Medical Research Institute of Infectious Diseases, Fort Detrick, USA). The expression plasmid pI18 for the GPC of ANDV strain CHI-7913 was kindly provided by Nicole Tischler (Molecular Virology Laboratory, Fundación Ciencia & Vida, Santiago, Chile) and has been described previously (Cifuentes-Munoz, Darlix et al. 2010). Expression plasmids encoding Lassa virus GPC strain Josiah and VSV-G have been reported previously (Torriani, Trofimenko et al. 2019). The expression vector encoding Ebola virus strain Makona was kindly provided by Mark Page (The National Institute for Biological Standards and Control, South Mimms, UK). Flag-hTim1 (Addgene plasmid # 49207) was a gift from L. Kane (de Souza, Oriss et al. 2005) and pLXSN-Axl (Addgene plasmid # 65222) a gift from A. Ullrich (Zhang, Knyazev et al. 2008). Human TIMD4 was obtained from Sino Biological Inc. Purified goat anti-human Axl IgG polyclonal antibody (pAb, #AF154), purified goat anti-human TIM-1/KIM-1/HACVR IgG pAb (#AF1750), phycoerythrin (PE)-conjugated donkey anti-goat IgG pAb, purified donkey anti-goat IgG pAb (#AF109) and monoclonal mouse IgG (#MAB002) were from R&D Systems. Purified rabbit anti-TIM4 pAb (#ab47637) was from Abcam. Purified mouse anti-human integrin αvβ3 monoclonal antibody (MAB1976) was from Millipore. PE-conjugated goat anti-mouse pAb and alexa 647-conjugated goat anti-rabbit pAb were from Invitrogen. The VSV neutralizing antibody I1 (I1 mAb) has been described (Holland, de la Torre et al. 1989).

The chemicals included G418 (Promega), R428 (Selleckem), duramycin (Chem Cru), heparan sulfate sodium salt from bovine kidney (H7640) (Sigma). Lipofectamine® 3000 Transfection Kit were obtained from Invitrogen. The CellTiter-Glo® Assay System and the ONE-Glo™ Luciferase Assay System were obtained from Promega (Madison WI).

### Cells

Human lung carcinoma alveolar epithelial (A549) cells, and human embryonic kidney (HEK293) cells were maintained in Dulbecco modified Eagle medium (DMEM)-10% [vol/vol] fetal calf serum (FCS) at 37°C under 5% CO_2_ atmosphere. Monkey kidney epithelial (Vero E6) cells were maintained in Biochrom minimum essential media (MEM) with Earle’s salts supplemented with 10% [vol/vol] FCS, 0.625% L-glutamine, 0.5% penicillin-streptomycin and 0.5% NEAA (Biochrom) at 37°C. Stable HEK293H cell lines expressing hTIM-1 were selected and maintained in 0.5 mg/ml G418. For infection studies, cells were seeded in 96-well plates at 35′000 cells/well and cultured 16 h until cell monolayer formation. Transient expression of hTIM-4 in HEK293H cells was obtained by using Lipofectamine® 3000 Transfection Kit (Invitrogen) following manufacturer’s instructions.

### Pseudotype virus production and viruses

The pseudotype viral system was based on the recombinant VSV*ΔG-Luc vector in which the glycoprotein gene (G) had been deleted and replaced with genes encoding green fluorescent protein (EGFP; indicated by an asterisk) and luciferase (Luc) (Berger Rentsch and Zimmer 2011). Pseudoviruses were generated as reported previously (Torriani, Mayor et al. 2019, Torriani, Trofimenko et al. 2019). Recombinant human adenovirus serotype 5 (AdV-5) expressing EGFP has been described previously (Oppliger, Torriani et al. 2016). HTNV strain 76/118 (Lee, Lee et al. 1978) was propagated in Vero E6 cells (Vero C1008; ATCC CLR 1586) in the BSL-3 containment laboratory at Spiez laboratory.

### Flow cytometry analysis

For cell surface staining, cells were first fixed with 2% paraformaldehyde (PFA) in PBS for 1 h at 4°C. The cells were then incubated with either polyclonal goat anti-hTIM-1, polyclonal goat anti-Axl, monoclonal mouse anti-αvβ3, or polyclonal rabbit anti-TIM-4 antibodies for 1 h at 4°C in PBS containing 1% [vol/vol] FCS and 0.1% [wt/vol] sodium azide (FACS buffer). Cells were washed twice in cold FACS buffer and incubated with the appropriate secondary antibody (PE-conjugated donkey anti-goat IgG, PE-conjugated goat anti-mouse IgG and alexa 647-conjugated goat anti-rabbit) for 45 min at 4°C in cold FACS buffer. Cells were finally washed twice and resuspended in PBS. Flow cytometry was performed using the BD Accuri^TM^ C6 (BD Bioscience) flow cytometer.

### Antibody perturbation assay

Confluent monolayers of A549 cells were washed twice with serum-free DMEM for 20 min at 37°C, and incubated for 1 h at 4°C in serum-free DMEM in presence of either goat polyclonal anti-hTIM-1, goat anti-Axl pAb, mouse monoclonal anti-αvβ3 antibodies or the isotype controls (anti-mouse or normal goat IgG) at 30 nM. Infection with the pseudoviruses was performed at 200 IU/well for 90 min at 37°C in the presence of the antibodies and 10% FCS. The inoculum was removed and cells were further incubated for 16 h at 37°C in DMEM, 10% FCS supplemented with 20 mM of ammonium chloride to prevent secondary infection. Infection was quantified by counting EGFP-positive infected cells per well using the FLoid Cell Imaging Station (ThermoFisher Scientific).

### Infection with pseudoviruses

Confluent monolayers of HEK293H hTIM-1/-4 cells were infected with the indicated viruses at different IU for 1 h at 37°C. Fresh DMEM containing 10% FCS was then added to the cells. After 16 h of incubation at 37°C in the presence of pseudoviruses, infection was quantified either by counting the number of EGFP-positive infected cells per well or by measuring luciferase with the TriStar LB 941 (Berthold Technologies) luminometer.

### Drug inhibition assay

Confluent monolayers of A549 cells were pre-treated with drugs for 30 min at 37°C under 5% CO_2_, followed by infection with the indicated viruses in presence of the drugs for 1.5 h at 37°C. After incubation, cells were washed twice with DMEM, 10% FCS supplemented with 20 mM of ammonium chloride and incubated for 16 h at 37°C with the presence of ammonium chloride. Finally, infection was quantified by counting EGFP-positive infected cells per well.

For time-of-addition experiments, 1 μM R428 was added at different time points: 45 minutes before viral infection (−0.5 h), during viral infection (0 h), and 1.5 h, 2.5 h and 4.5 h after viral infection (+1.5 h, +2.5 h, +4.5 h). Infection with the indicated viruses was performed in presence of the drug for 1.5 h at 37°C under 5% CO_2_. The cells were subsequently washed twice with DMEM, 10% FCS supplemented with 20 mM of ammonium chloride and incubated for 16 h at 37°C with the presence of ammonium chloride and the drug. Productive infection was quantified by counting EGFP-positive infected cells per well.

### Treatment of viral particles with duramycin or heparan sulfate

Confluent monolayers of A549 cells were grown 16 h at 37°C with DMEM 10% FCS in a 96-well plate. Indicated pseudoviruses (200 IU/well) were incubated with duramycin or soluble heparan sulfate for 1 h at RT and added to cells. After 90 min of infection, inoculum was removed and fresh medium containing 10% FCS and 20 mM ammonium chloride was added. Infection was quantified 16 h after by counting the number of EGFP-positive infected cells per well.

### Infection with authentic infectious HTNV, RNA isolation and RT-qPCR

HTNV (strain 76/118) was pre-treated for 30 min at RT with increasing concentrations of duramycin. Vero E6 cells were subsequently infected for 1 h at 37°C in presence of the drug, washed once with MEM 2% and incubated with MEM 2% containing appropriate duramycin concentrations for 5 days. Viral RNA was isolated using EZ1 Virus Mini Kit v2.0 (QIAGEN). Quantification of viral RNA was performed using a real-time RT-PCR assay specific for HTNV nucleocapsid coding region in a LightCycler® 96 (Roche) following manufacturer’s instructions.

### Statistical analysis

Graphical representation and statistical analysis were performed using GraphPad Prism 7 software. Data are means + SD (n=3) with p-value *: p ≤ 0.05; **: p ≤ 0.01; ***: p ≤ 0.001. P values of <0.05 were considered statistically significant.

## ACKNOWLEDGEMENTS

The authors thank Professor Stefan Kunz for helpful discussions and support. We thank the virology team at Spiez Laboratory for helping to establish the Hantaan virus culture and validation experiment. We also thank Dr. Nicole Tischler (Molecular Virology Laboratory, Fundación Ciencia & Vida, Santiago, Chile) for the expression plasmid pI18 GPC ANDV. We further thank Dr. Connie Schmaljohn (USAMRIID, Fort Detrick, MD, USA) for the plasmid pWRG/HTNV-M, and Mark Page (The National Institute for Biological Standards and Control, South Mimms, UK) for the expression plasmid pCAGGS-GUI14. This research was supported by Swiss Federal Office for Civil Protection (Grants Nr. 353006233/Stm 353008560/Stm and 35 3008553 to O.E. and S.K.), Swiss Lung Association (Grant Nr.: 2016-06_Kunz to SK, OE and SR). JM was supported by funds from the University of Lausanne to Stefan Kunz.

